# Overexpression of tomato *SlBBX16* and *SlBBX17* impacts fruit development and gibberellin metabolism

**DOI:** 10.1101/2023.08.12.553068

**Authors:** Valentina Dusi, Federica Pennisi, Daniela Fortini, Alejandro Atarés, Stephan Wenkel, Barbara Molesini, Tiziana Pandolfini

## Abstract

BBXs are B-Box zinc finger proteins that can act as transcription factors and regulators of protein complexes. Several BBX proteins play important roles in plant development. Two *Arabidopsis thaliana* microProteins belonging to the BBX family, named miP1a and miP1b, homotypically interact with and modulate the activity of other BBX proteins, including CONSTANS, which transcriptionally activates the florigen, *FLOWERING LOCUS T*. In tomato, the closest homologs of miP1a and miP1b are the microProteins *Sl*BBX16 and *Sl*BBX17. To deepen our understanding of the role of tomato microProteins in flowering, we constitutively expressed *Sl*BBX16/17 in *Arabidopsis* and tomato and examined possible interacting partners. Overexpression of the two tomato microProteins in *Arabidopsis* caused a delay in the flowering transition; however, the effect was weaker than that observed in *Arabidopsis* plants overexpressing the native miP1a/b. In tomato, overexpression of *SlBBX17* prolonged the flowering period; this effect was accompanied by downregulation of the flowering inhibitors *Self Pruning* (*SP*) and *SP5G*. *Sl*BBX16 and *Sl*BBX17 are able to hetero-oligomerize with TCMP-2, a cystine-knot peptide involved in flowering pattern and fruit development in tomato. Increasing the expression of both tomato microProteins also caused alterations in fruit development: overexpression of *SlBBX17* resulted in a diminished number and size of ripe fruits as compared to WT plants, while overexpression of *SlBBX16* caused delayed fruit production up to the breaker stage. These effects were associated with changes in the expression of genes regulating gibberellin content.

## Introduction

Crop productivity is the result of the interaction between the plant genetic background, the environmental cues and the agronomic practices. In the face of climate change, it becomes increasingly important to match the phenology of crop cultivars with the environmental conditions to maintain high yields. Therefore, the availability of cultivars with different phenological requirements could be advantageous for optimising plant reproduction and productivity. In horticultural plants, flowering, fruit set and the onset of ripening are processes markedly affected by temperature and photoperiod, and regulated by internal signals such as hormones, transcription factors and adaptor molecules. Among the different regulatory proteins known to control reproductive development, members of the B-Box (BBX) protein family have emerged as important molecular players that integrate environmental cues, for instance, light and temperature, with endogenous signaling pathways (Gangappa and Botto 2014; Lira et al., 2020; Yadav et al., 2020).

The BBX family represents a group of zinc (Zn)-finger proteins that are involved not only in reproductive development, but also in many other physiological processes, such as photomorphogenesis (Crocco et al.,2010; Holtan et al., 2011; Fan et al., 2012), anthocyanin accumulation, seed germination, carotenoid biosynthesis, and responses to biotic and abiotic stresses (Wang et al., 2013; Kiełbowicz-Matuk et al., 2014; Vaishak et al., 2019; Xu et al., 2022). The BBX family is characterized by the presence of one or two Zn-finger-containing BBX domain(s) in the N-terminal region (Khanna et al., 2009; Gangappa and Botto, 2014; Graeff et al.,2016). Previous studies suggested that the B-Box domain plays a crucial role in protein-protein interactions. Some BBX proteins may possess a CCT domain, which is associated with a role in transcriptional regulation and nuclear transport (Gendron et al., 2012; Crocco and Botto, 2013). CCT-containing BBXs are likely transcription factors, while BBXs lacking CCT domain might have other functions, linked to their protein-protein interaction capacity. *Arabidopsis* BBX proteins have been classified into five subgroups according to different combinations of the above-mentioned domains. Members of group I are characterized by the presence of B1, B2, and CCT domains, as are members of group II; however, some differences have been observed between the two groups in the B2 consensus sequences (Crocco and Botto, 2013; Gangappa and Botto, 2014). The presence of B1 and CCT, B1 and B2, and a single B1 domain characterizes members of group III, IV, and V, respectively (Khanna et al., 2009; Gangappa and Botto, 2014;). Similarly, following structural features, BBX proteins from other species such as tomato, potato, rice, and maize have been grouped into subfamilies (Huang et al., 2012; Chu et al., 2016; Li et al., 2017; Talar et al., 2017).

A. *thaliana* CONSTANS (*At*CO), one of the first BBX proteins to be identified and characterized, belongs to subgroup I. *At*CO plays a crucial role in photoperiodic control of flowering time, enabling the transcriptional activation of the *FLOWERING LOCUS T* (*FT*) in response to long day (LD) conditions. Proteins homologous to *At*CO have been shown to contribute to flowering regulation in other species also with different photoperiodic requirements (Doi et al., 2004; Miller et al., 2008; Meng et al., 2011; Campoli et al., 2012; Kikuchi et al., 2012; Yang et al., 2014; Ping et al., 2019; Wang et al., 2019).

Other members of the *Arabidopsis* BBX family were successively discovered to be implicated in flowering control (Cheng and Wang, 2005; Park et al., 2011; Li et al., 2014; Tripathi et al., 2017). Recently, two *Arabidopsis* BBX proteins of the group V, microProtein (miP) miP1a and miP1b, (also referred to as *At*BBX31 and *At*BBX30, respectively) were shown to modulate *At*CO activity (Graeff et al., 2016; Rodrigues et al., 2021). MiP1a and miP1b can interact with both *At*CO and TOPLESS (TPL), leading to the formation of a trimeric complex that limits *At*CO-mediated induction of *FT* expression. Consistently, *Arabidopsis* plants overexpressing miP1a and miP1b, grown under LD conditions, showed delayed flowering (Graeff et al., 2016). In tomato, the microProteins *Sl*BBX16 and *Sl*BBX17 are the closest homologs of miP1a and miP1b, but their role in tomato reproductive development is largely elusive. The features of group V BBXs - a single B-Box domain and the lack of CCT - suggest they act as microProteins regulating other larger multidomain complexes at the post-translational level (Eguen et al., 2015).

In this regard, *Sl*BBX16 was found to interact with the tomato cystine-knot peptide 2 (TCMP-2) (Molesini et al., 2020). TCMP-2 is specifically expressed in reproductive organs, its expression is low in pre-anthesis flower buds and gradually increases after fertilization, reaching a maximum in green and ripe fruits (Pear et al., 1989; Cavallini et al., 2011; Treggiari et al., 2015). Increased TCMP-2 expression in pre-anthesis flower buds has been demonstrated to lead to altered flowering pattern as well as early fruit production and a slight delay in the initiation of ripening (Molesini et al., 2018; Molesini et al., 2020).

In tomato, which is a day-neutral species, the flowering transition is principally regulated by the balance between the activity of the florigen *Single Flower Truss* (*SFT*), which is the ortholog of *FT*, and that of antiflorigens, such as *Self Pruning* (*SP*) and *SP5G*, which maintain vegetative growth. Recently, it has been shown that *Sl*COL1, the ortholog of *At*CO, is able to bind the promoter region of *SFT* to repress its expression (Cui et al., 2022). Accordingly, RNA silencing of *SlCOL1* led to the promotion of flowering and increased fruit yield.

Our work was aimed at elucidating the functional role of *Sl*BBX16 and *Sl*BBX17 in tomato reproductive development. We demonstrated that *SlBBX17* overexpressing (*Sl*BBX17OE) plants show a prolonged period of flowering associated with a reduced expression of the *SP* and *SP5G* flowering inhibitors. Furthermore, we observed a delay in the early phases of fruit growth in *Sl*BBX16OE plants and changes in ripening in *Sl*BBX17OE plants,. The overexpression of both microProteins induced modifications in the expression pattern of genes regulating gibberellin (GA) metabolism.

## Results

### Ectopic overexpression of *SlBBX16* and *SlBBX17* influences the flowering transition in *Arabidopsis*

The microProteins *Sl*BBX16 and *Sl*BBX17 show high sequence similarity to miP1a and miP1b (Figure S1A), which control flowering in *Arabidopsis* and, when overexpressed, delay flowering (Graeff et al., 2016). To test whether *Sl*BBX16 and *Sl*BBX17 can exert a similar effect, we analysed the flowering behaviour of *Arabidopsis Sl*BBX16OE and *Sl*BBX17OE plants compared to Col-0 WT (Figure 1 and Figure S1B, C). In the *Sl*BBX16OE plants, the number of rosette leaves at bolting did not differ from the control plants (Figure 1A). Also, the days from sowing to flowering did not vary (Figure 1B). On the other hand, both *Sl*BBX17OE lines displayed an increased number of rosette leaves at bolting and a longer time to reach flowering under LD photoperiodic conditions (Figure 1C, D). The observed inhibitory effect of *Sl*BBX17 on flowering was less pronounced than that produced by overexpressing miP1a and miP1b in *Arabidopsis* (Graeff et al., 2016). To examine whether *Sl*BBX16 and *Sl*BBX17 can substitute the function of miP1a/b, we overexpressed them also in *Arabidopsis* miP1a/b double KO mutant, which displays earlier flowering than WT (Heng et al., 2019). In both *Sl*BBX16OE lines and in the *Sl*BBX17OE #5, the number of leaves at bolting (Figure 1E, G) increased significantly (*Sl*BBX16OE #2 13.3 ± 1.5; #3 13.6 ±1.9; *Sl*BBX17OE #5 12.7 ± 1.6) compared to miP1a/b double KO mutant (11.7± 1.2) and was slightly lower than or similar to Col-0 WT (13.6 ±0.3). The number of days to bolting was increased significantly in *Sl*BBX16OE #2 (31.7± 1.6) and in both *Sl*BBX17OE lines (33.2 ± 3.3 and 34.4 ± 3.1 days for #1 and #5, respectively) (Figure 1F, H) reaching values similar to those observed in Col-0 WT (Fig. 1B, D). These results suggest that *Sl*BBX16 and *Sl*BBX17 may impact the flowering transition in *Arabidopsis* and partially rescue the function of endogenous miP1a/b. Since the flowering delay exhibited by *Sl*BBX16OE and *Sl*BBX17OE plants resembles, albeit in an attenuated form, the phenotype shown by miP1a/b-overexpressing plants (Graeff et al.,2016), we examined via Y2H whether *Sl*BBX16 and *Sl*BBX17 interact with *At*CO as already demonstrated for miP1a/b (Graeff et al., 2016). Under our experimental conditions, we did not observe a direct interaction between *Sl*BBX16 and *Sl*BBX17 and *At*CO (Figure S2).

**Figure 1.**
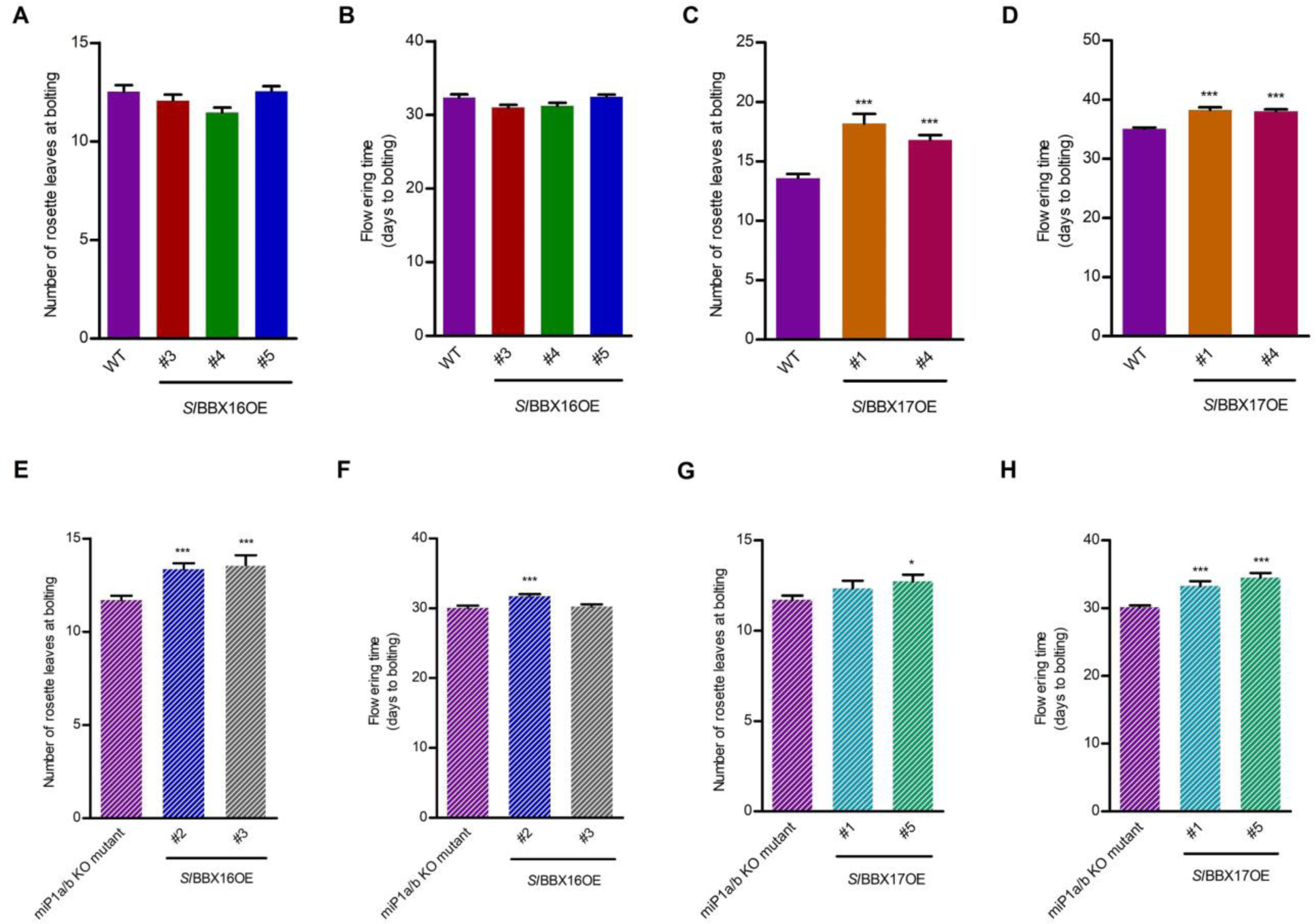
Overexpression of *SlBBX16* and *SlBBX17* in *Arabidopsis* WT (Col-0) and miP1a/b KO mutant. The transition from vegetative to reproductive development was determined by counting the number of rosette leaves at the bolting stage and the number of days to bolting. **(A, B)** Three lines overexpressing *SlBBX16* (*Sl*BBX16OE #3, #4, and #5) and **(C, D)** two lines overexpressing *SlBBX17* (*Sl*BBX17OE #1, and #4) were compared with WT. Values are mean ± SE (n= 19-27, for panels A and B; n=16-26, for panels C and D). **(E, F)** Two *Sl*BBX16OE lines (#2, and #3) and **(G, H)** two *Sl*BBX17OE lines (#1, and #5) were compared with miP1a/b double KO mutant. Values are mean ± SE (n= 13-26, for panels E and F; n= 18-26 for panels G and H). *P<0.05 and ***P < 0.001 versus the respective control (Student’s t-test).

### Overexpression of *SlBBX16* alters fruit set and delays fruit growth in MicroTom

We recently demonstrated that *Sl*BBX16 interacts with TCMP-2, a tomato cystine-knot metallocarboxypeptidase inhibitor (Molesini et al., 2020). TCMP-2 is specifically expressed in reproductive organs (Cavallini et al., 2011; Molesini et al., 2018) and, when overexpressed in flower buds, caused alteration in flowering pattern and early fruit setting in the tomato cultivar UC82 (Molesini et al., 2018; Molesini et al. 2020). When *TCMP-2* is globally overexpressed (i.e., using the *CaMV 35S* promoter) in the cultivar MicroTom, we have observed an anticipated formation of the primary inflorescence and a reduction in plant height, suggesting an accentuated determinate habit (Figure S3). The fruit set calculated on the first three inflorescences did not change significantly (data not shown). We then analyzed the role of *Sl*BBX16 microProtein in reproductive development by overexpression in MicroTom. The three overexpressing lines #3, #4, and #20 (Figure S4A) showed no changes in the number of leaves before the first inflorescence compared to the WT plants (Figure 2A), while the shoot height at the first inflorescence was significantly reduced only in the *Sl*BBX16OE #3 (Figure 2B). These data suggest that flowering transition was not greatly affected by the *SlBBX16* overexpression. In addition, the number of flowers measured in the first three inflorescences was comparable in the WT and *Sl*BBX16OE lines (Figure 2C). However, fruit set, calculated as a percentage of number of fruits over number of flowers in the first three inflorescences, was significantly lower in the *Sl*BBX16OE lines than in the WT plants (Figure 2D, E). We have also observed a delay in the initial fruit formation and in the onset of maturation assessed as number of fruits at the green stage (Figure 3A) and number of fruits at the breaker stage or in maturation (Figure 3B) overtime, respectively. Considering the total fruit production harvested at 110 days after sowing, in the *Sl*BBX16OE lines there was a partial compensation of the fruit set impairment in the first three inflorescences (Figure 3C). The weight of WT and transgenic fruits was similar (Figure 3D), as was the percentage of red and green fruits (data not shown).

**Figure 2.**
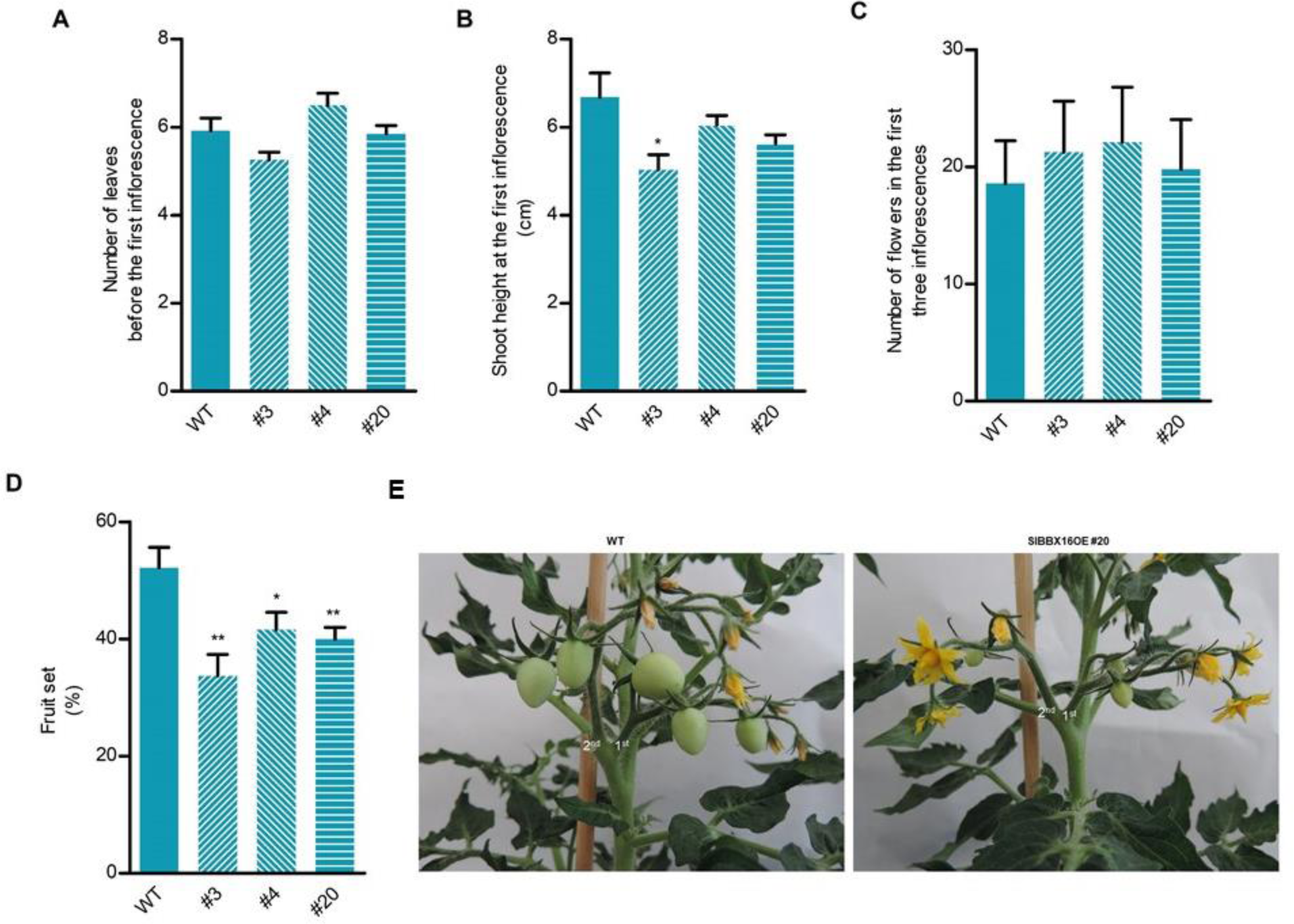
Transition to flowering and fruit set parameters of *Sl*BBX16OE plants. **(A)** Number of leaves before the first inflorescence **(B)** Shoot height at the first inflorescence. **(C)** Number of flowers in the first three inflorescences. **(D)** Fruit set of the first three inflorescences calculated as the percentage of the number of fruits over the number of flowers. (**E**) Representative pictures of the first two inflorescences of WT and *Sl*BBX16OE #20 plants of the same age. The values reported are means ± SE (n= 10-13). Student’s t-test was used for the statistical analysis (*P < 0.05; **P < 0.01).

**Figure 3.**
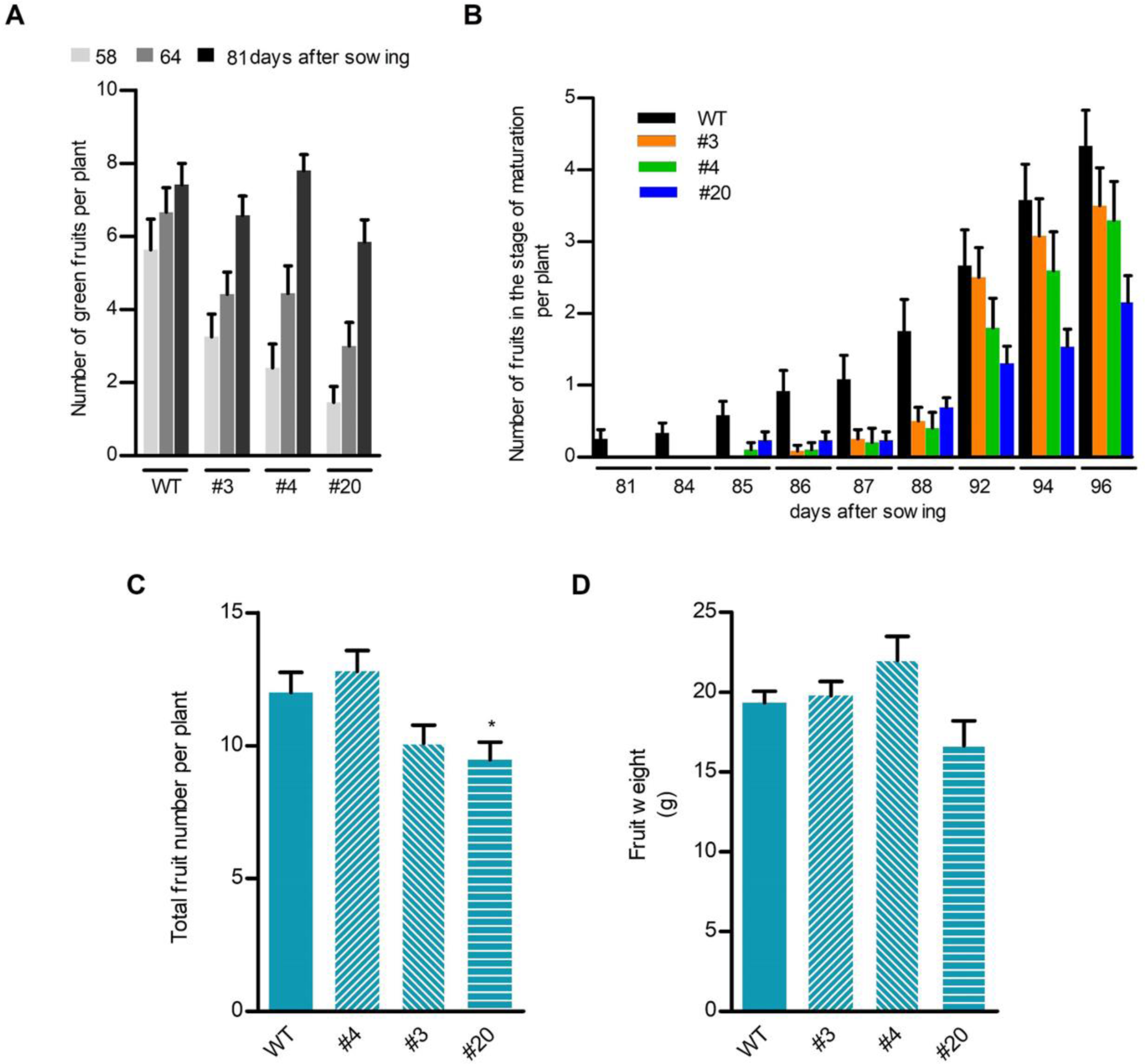
Fruit growth and production in *Sl*BBX16OE plants. **(A)** Number of green fruits recorded from 58 to 81 days after sowing. **(B)** Number of fruits in the stage of maturation. **(C)** Total fruit number per plant recorded 110 days after sowing. **(D)** Average fruit weight. The values reported are means ± SE (n= 10-13). Student’s t-test was used for the statistical analysis (*P < 0.05).

### Overexpression of *SlBBX17* prolongs flowering and reduces ripe fruit production in MicroTom

The microProteins *Sl*BBX16 and *Sl*BBX17 are highly homologous, with 46% identity (Figure S1A). The two genes are both expressed in the ovary and young fruit and are co-expressed with *TCMP-2* (Figure S4B and S5A, Cavallini et al., 2011, Molesini et al., 2020). Furthermore, Y2H analysis suggests that *Sl*BBX17 and *Sl*BBX16 share a common interacting partner, TCMP-2 (Figure S5B and Molesini et al. 2020).

We investigated the phenotypic consequences of *SlBBX17* overexpression in MicroTom by comparing two independent *Sl*BBX17OE transgenic lines (#15, and #18) to WT plants (Figure S5C). Differently from what was previously reported by overexpressing *SlBBX17* in the indeterminate cultivar Ailsa Craig (Xu et al., 2022), we did not observe retardation in vegetative growth and a reduction in leaf size (data not shown). However, because of the stunted growth of MicroTom it is likely that these growth effects were masked by other mutations. In agreement with Xu and collaborators (2022), *Sl*BBX17OE plants showed moderate resistance to high temperatures (Figure S5D). When we monitored the transition from vegetative to reproductive development in *Sl*BBX17OE plants, we observed a reduction in the height of the shoot bearing the primary inflorescence compared to WT plants, but no changes in the number of leaves before the first inflorescence (Figure 4A, B). The number of flowers in the first three inflorescences was similar in the WT and transgenic lines (Figure 4C), but when recording the number of flowers at anthesis over time, we noticed a different trend in the WT and transgenic plants (Figure 4D, E). In the WT plants, the number of open flowers increased from 39 to 47 days after sowing (das) but decreased sharply thereafter. In the transgenic plants, we observed a tendency to persist in producing flowers over time: in line #15 the number of flowers was higher than in WT from 61 das onwards and in line #18 this effect was detected from 75 das onwards. Given the differences in the duration of flowering between WT and *Sl*BBX17OE plants (Figure 4D, E), we analysed the expression of *SFT*, the ortholog of *FT*, and the flowering inhibitors, *SP* and *SP5G*. The dynamic ratio between florigen and antiflorigens regulates shoot termination in tomato, which is associated with flowering (Lifschitz et al., 2006; Lifschitz et al., 2014; Cao et al., 2016; Rajendran et al., 2021). The expression of *SFT* did not vary, whereas the expression of *SP* and *SP5G* was reduced in the transgenic plants compared to WT (Figure 4F-H). This suggests that the prolonged flowering observed in *Sl*BBX17OE might be associated with a weaker antagonistic effect of SP on SFT. Tomato is a day-neutral plant and the mechanism controlling the flowering process in this species is not fully elucidated. In this regard, the biological functions of CO-like (COL) genes in tomato remain elusive. Recently, *Sl*COL1 (Solyc02g089540) was shown to interact with the *SFT* promoter repressing its transcription and consequently inhibiting the flowering transition (Cui et al., 2022). Considering that miP1a/b interacts with *At*CO regulating its activity, we tested by Y2H the interaction between *Sl*BBX16 and *Sl*BBX17 and *Sl*COL1. The absence of interaction between tomato BBX proteins and *Sl*COL1, would suggest that either *Sl*BBX16 and *Sl*BBX17 do not bind tightly to *Sl*COL1 under our experimental conditions or that their interaction requires additional factors that are absent in yeast (Figure S2).

**Figure 4.**
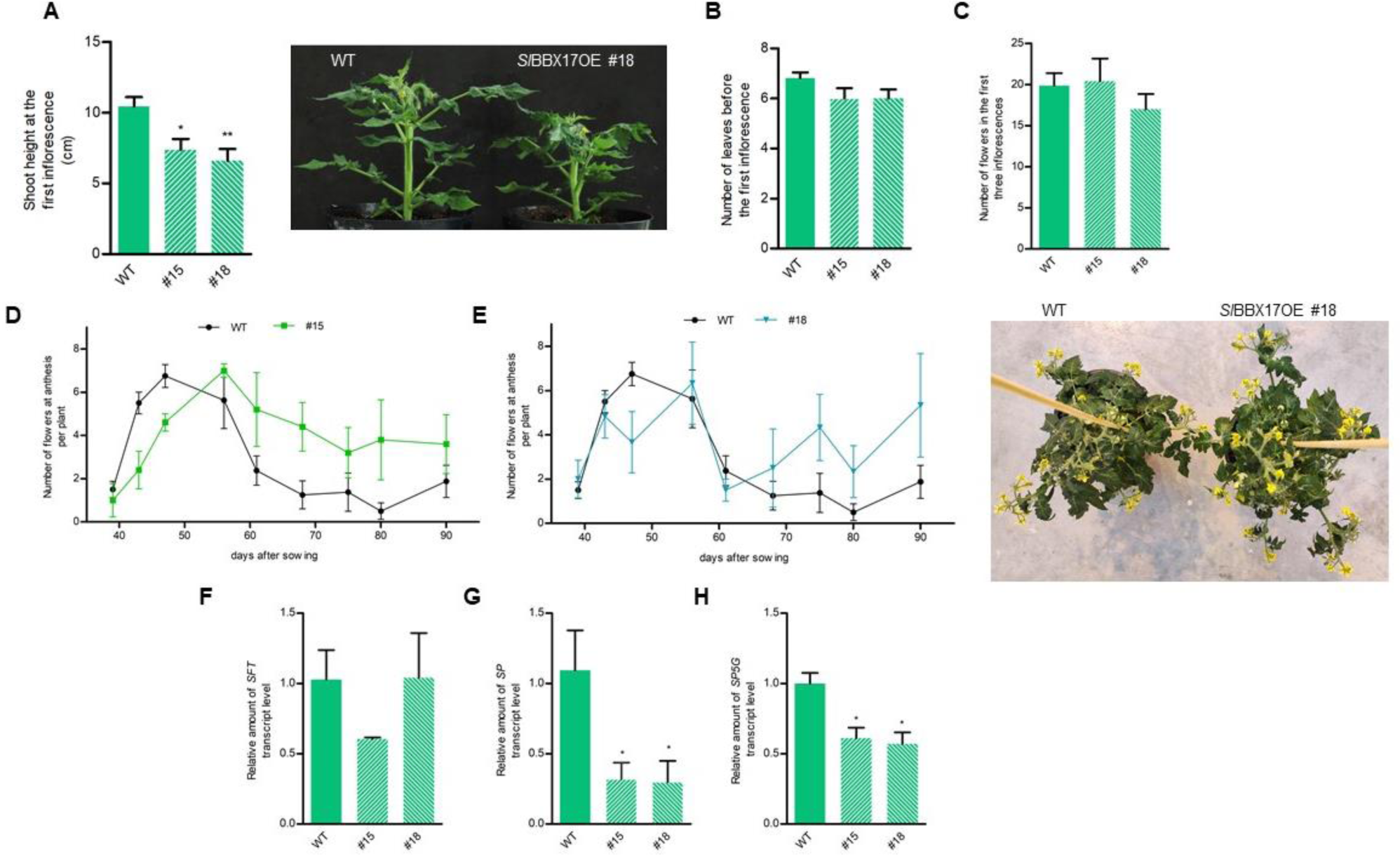
Fruit growth and production in *Sl*BBX17OE plants. **(A)** Shoot height at the first inflorescence. **(B)** Number of leaves before the first inflorescence. **(C)** Number of flowers in the first three inflorescences. **(D** and **E)** Number of flowers at anthesis per plant from 39 to 90 days after sowing. The values reported are means ± SE (n= 5-8). Phenotypical aspect of WT and *Sl*BBX17OE #18 at 85 days after sowing (panel E, on the right). **(F-H)** Expression level of *SFT*, *SP,* and *SP5G* of WT and *Sl*BBX17OE plants. Values are means ± SE of three biological replicates. Student’s t-test was used for the statistical analysis (*P < 0.05; **P < 0.01).

We then evaluated the fruiting process in the *Sl*BBX17OE lines assessing fruit set and fruit productivity. The fruit set recorded in the first three inflorescences and the total number of fruits per plant evaluated 4 months after sowing were similar to WT plants (Figure 5A, B). However, the proportion of ripe fruits out of the total number of fruits in the transgenic lines was reduced (Figure 5C, F). In addition, ripe fruits were smaller and contained fewer seeds (Figure 5D, E). We have also observed a different weight distribution of green fruits with a decreasing trend in *Sl*BBX17OE fruits (Figure S6).

**Figure 5.**
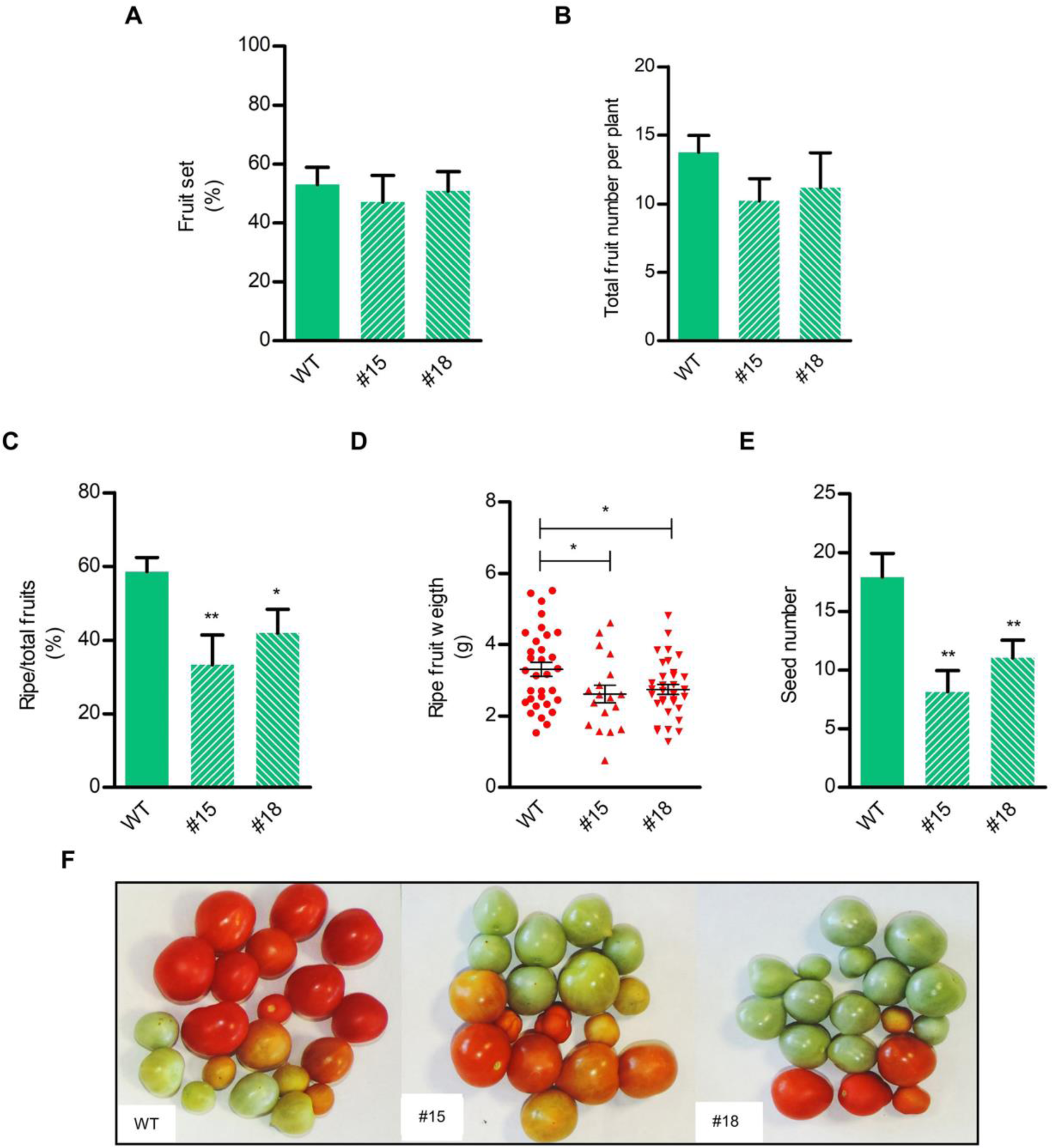
Fruit growth and production in *Sl*BBX17OE plants. **(A)** Fruit set percentage of the first three inflorescences calculated as the percentage of number of fruits over number of flowers. **(B)** Total fruit number per plant, **(C)** percentage of ripe over total fruits collected 110 days after sowing **(D)** weight of ripe fruits and **(E)** number of seeds collected from ripe fruits. **(F)** Representative picture of fruit production. The values reported are means ± SE (n 8-14 for panel A, B and C; n=18-34 for panel D and E). Student’s t-test was used for the statistical analysis (*P < 0.05; **P < 0.01).

### Overexpression of *SlBBX16* and *SlBBX17* leads to altered expression of genes involved in GA metabolism

To elucidate if *Sl*BBX16 and *Sl*BBX17 affect the hormone homeostasis that controls fruit development, we examined in *Sl*BBX16OE and *Sl*BBX17OE immature green fruits, a phase of active growth and preparation for ripening, the expression of a set of genes implicated in ethylene, abscisic acid (ABA), and gibberellin (GA) metabolism. Aminocyclopropane-1-carboxylic acid synthase 1A (ACS1A) and ACS6 encode enzymes of the ethylene biosynthetic system 1 that triggers the transition to the breaker stage in green fruit (Barry et al., 2000; Alexander and Grierson, 2002). Their expression did not vary in *Sl*BBX16OE and *Sl*BBX17OE fruits (Figure 6A, B, E, F), suggesting that ethylene biosynthesis at this stage is similar to that in WT fruits. To test a possible alteration of ABA metabolism, we analysed the expression of 9-cis-epoxycarotenoid dioxygenase (*SlNCED-1*), the key determinant of ABA synthesis, and *SlZFP2*, a zinc finger transcription factor involved in the crosstalk between ABA and ethylene in the regulation of fruit ripening in tomato (Weng et al., 2015). The expression of *SlNCED-1* increased in two (#3 and #4) out of the three *Sl*BBX16OE lines, while it remained unaltered in *Sl*BBX17OE fruits compared to WT (Figure 6D, H). The transcript level of *SlZFP2* was unchanged in *Sl*BBX16OE and *Sl*BBX17OE fruits (Figure 6C, G). A reduced expression of the key GA biosynthetic gene, GA20 oxidase 2 (*SlGA20ox2*), was recorded in the three *Sl*BBX16OE lines and in the *Sl*BBX17OE #15 (Figure 7B, D). On the other hand, in the SlBBX17OE #18, *SlGA2ox4*, a gene encoding an enzyme responsible for GA catabolism, was induced (Figure 7C). Genes involved in GA metabolism are regulated by a negative feedback loop; high GA levels or signals inhibit the GA biosynthetic gene GA20ox2 and induce GA deactivation genes, like GA2ox4 (Livne et al., 2015). The upregulation of GA2ox4 observed in *Sl*BBX17OE young fruit correlates with a higher content of bioactive GAs, GA1 and GA4 (Figure 7E). In *Sl*BBX16OE fruits, the downregulation of *GA20ox2* is not accompanied by alteration in GA1 and GA4 content (Figure 7E). These data suggest that the phenotypes observed in *Sl*BBX16OE and *Sl*BBX17OE fruits might be due to altered GA metabolism. Interestingly, treatment with exogenous GA induced the expression of both *Sl*BBX16 and *Sl*BBX17 (Figure 7F, G).

**Figure 6.**
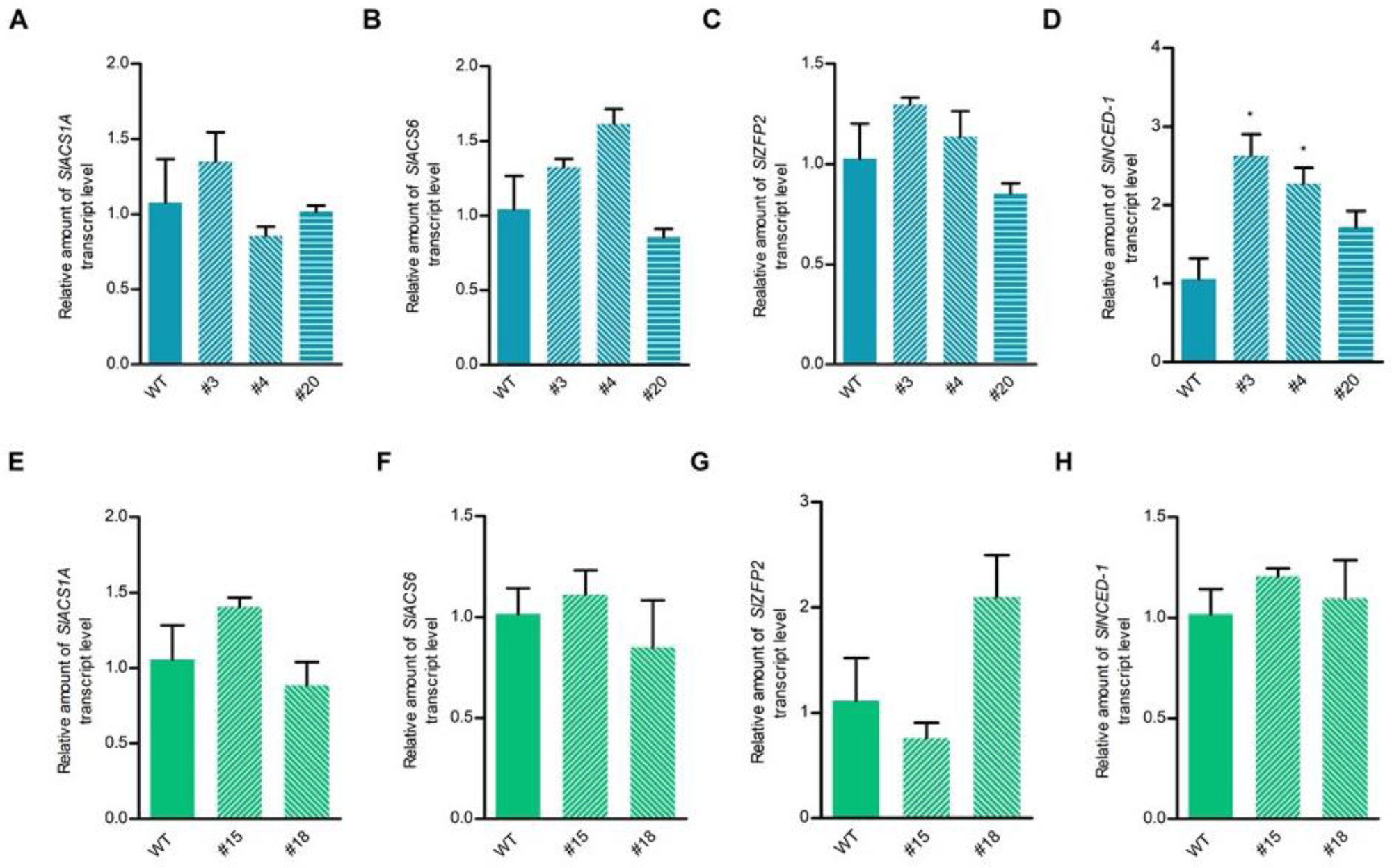
Expression of genes implicated in ethylene and abscisic acid metabolism. Transcript levels of *ACS1A*, *ACS6*, *ZFP2*, and *NCED-1* in WT and *Sl*BBX16OE **(A-D)** and *Sl*BBX17OE **(E-H)** green fruits. Values are means ± SE of three biological replicates. Student’s t-test was used for the statistical analysis (*P < 0.05; **P < 0.01).

**Figure 7.**
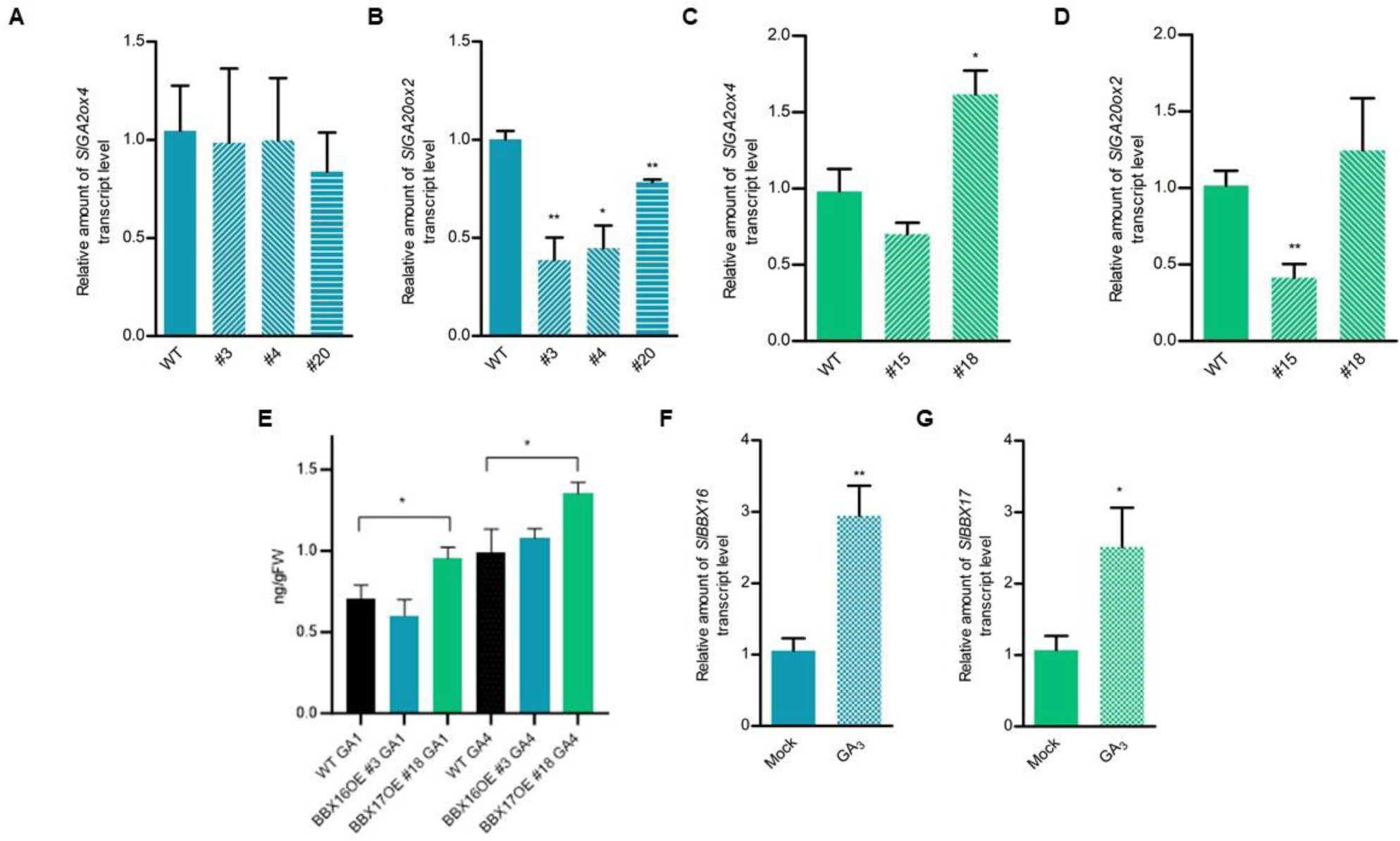
GA metabolism alteration in *Sl*BBX16OE and *Sl*BBX17OE lines. Expression analysis of genes implicated in GA catabolism (*GA2ox4*) and biosynthesis (*GA20ox2*) in WT and *Sl*BBX16OE **(A, B)** and *Sl*BBX17OE **(C, D)** fruits. **(E)** Bioactive GAs (GA1 and GA4) content in immature green fruits of *Sl*BBX16OE #3 and *Sl*BBX17OE #18 in comparison with WT. Response of *Sl*BBX16 **(F)** and *Sl*BBX17 **(G)** to GA. Relative expression of *SlBBX16* and *SlBBX17* was evaluated in shoots of seedlings treated for 24 h with 5 µM GA_3_ in comparison with mock-treated ones. Values are mean ± SE of three biological replicates. Student’s t-test was used for the statistical analysis (*P < 0.05; **P < 0.01).

## Discussion

Plant microProteins are small polypeptides characterized by a single domain, involved in protein-protein interaction. From an evolutionary point of view, miPs can originate from larger proteins by duplication and domain loss (*trans-*miPs) or derive from alternative splicing or alternative transcription start and stop sites (*cis*-miPs) (Eguen et al., 2015; Kushwaha et al., 2022). Their capacity to interact with homologous proteins of larger size (homotypic interaction) is at the basis of their activity as posttranslational regulators of protein complexes. Besides homotypic, plant miPs can engage also in heterotypic interactions with evolutionary unrelated proteins, widening the regulatory role of these proteins (Bhati et al., 2021).

*Arabidopsis* miP1a and miP1b are microProteins belonging to subclass V of the BBX family. They play a role in photomorphogenic development and control of flowering time. MiP1a and miP1b participate in seedling growth arrest after germination under stress conditions by interacting heterotypically with ABA-insensitive 5 (ABI5) and stabilizing it (Singh and Datta, 2023). During the transition from dark to light conditions, miP1a and miP1b inhibit the oligomerization of PHYTOCHROME-INTERACTING FACTORs (PIFs) and ETHYLENE-INSENSITIVE 3 (EIN3) (Wu et al., 2020). A homotypic interaction between miP1a/b and *At*CO regulates florigen *FT* transcription; the strong delay in flowering observed by overexpressing miP1a/b in *Arabidopsis* is caused by the recruitment of TOPLESS to miP1a/b-CO complex resulting in inhibition of *FT* transcription (Graeff et al., 2016).

In tomato, *Sl*BBX16 and *Sl*BBX17 microProteins are the closest homologs of miP1a/b. A study by Xu and collaborators (2022), reported that *SlBBX17* overexpression in an indeterminate tomato cultivar (i.e., Ailsa Craig), resulted in increased heat tolerance and growth retardation (Xu et al., 2022). More recently, *SlBBX17* was reported to participate also in the response to cold stress (Song et al., 2023). The effects of *SlBBX17* overexpression on reproductive development were not investigated in those studies. The first hints on the implication of *Sl*BBX17 and *Sl*BBX16 in reproductive development derive from the observation that *Sl*BBX16 and *Sl*BBX17 undergo heterotypic interactions with a tomato cystine-knot peptide, TCMP-2, which plays a role in both flowering pattern and fruit development (this work and Molesini et al., 2020). In the tomato cultivar UC82, overexpression of *TCMP-2* in flower buds promoted the termination of sympodial units (Molesini et al., 2020) and global overexpression in MicroTom resulted in the early appearance of the first inflorescence (this work). In addition, ectopic overexpression of *TCMP-2* anticipated flowering in *Arabidopsis* and in both tomato and *Arabidopsis*, TCMP-2 induced the transcription of florigen (Molesini et al., 2020). To investigate if TCMP-2 interacting partners play a role in flowering, *SlBBX16* and *SlBBX17* were overexpressed in *Arabidopsis* WT Col-0 and miP1a/b double KO mutant and in tomato. *Arabidopsis Sl*BBX16OE and *Sl*BBX17OE plants showed a weak delay in flowering, which is evident for *Sl*BBX16 only when overexpressed in a background depleted in miP1a/b function. This observation suggests that the mechanism of action of *Sl*BBX16 differs somewhat from that of *Sl*BBX17. The incomplete functional redundancy between the two proteins is confirmed by the fact that overexpression of *Sl*BBX16 in tomato did not cause appreciable alterations in flowering, whereas overexpression of *Sl*BBX17 resulted in protracted flower production. This latter effect is caused by a reduced expression of the antiflorigens, *SP* and *SP5G*. Thus, overexpression of *TCMP-2* and its partners produced contrasting effects on flowering, suggesting an antagonistic interaction. The evidence that both TCMP-2 and *Sl*BBX16/17 showed no direct interaction by Y2H with *At*CO and *Sl*COL1 (this work and Molesini et al., 2020) highlights the differences in the flowering control system between *Arabidopsis* and tomato. In this regard, *Sl*COL1 has been shown to bind the *SFT* promoter and negatively regulate its expression (Cui et al., 2022). Based on our results, TCMP-2 and *Sl*BBX17 activity in the tomato flowering process appeared independent of the *Sl*COL1/SFT regulatory module. We cannot exclude that *Sl*BBX17 can interact with other members of the *Sl*COL family.

This study demonstrates that *Sl*BBX16 and *Sl*BBX17 also participate in the regulation of fruit growth and maturation. In a recent paper, Lira and collaborators evaluated the transcript profiles of *SlBBX16* and *SlBBX17* in tomato fruits at different stages of development, starting from immature green up to 5 days after the breaker stage (Lira et al., 2020). During the stages of green fruit growth, the expression of *SlBBX16* progressively decreased (Lira et al., 2020). This expression pattern might indicate that during the period of fruit enlargement, *Sl*BBX16 activity should remain low, an observation consistent with the phenotype of *Sl*BBX16OE plants, in which fruit growth is delayed from the early green phases until the breaker stage. It is plausible that *Sl*BBX16 is one of the factors that restricts ovary expansion; therefore, its progressively reduced expression would be necessary for optimal fruit growth. The reduced fruit set observed in the first three inflorescences of *Sl*BBX16OE plants would support this hypothesis. The expression of *SlBBX17* is not sharply modulated during fruit development as for *SlBBX16* (Lira et al., 2020); its transcript level increases weakly from immature green to the breaker stage and then declines shortly after (Lira et al., 2020). In the present work, the primary effect of *SlBBX17* overexpression was observed at the ripening stage. The percentage of ripe to total fruits was markedly reduced, suggesting that the excess of this microProtein at the maturation phase hinders this process. The reduced weight of ripe *Sl*BBX17OE fruits and the similar tendency observed for green fruits indicate that *Sl*BBX17 is also involved in fruit enlargement. Although the two microProteins seem to contribute to regulating fruit development at different stages, they both lead in young fruits, when overexpressed, to modification in GA metabolism with either inhibition of GA biosynthetic genes or induction of GA deactivation gene. Remarkably, both *SlBBX16* and *SlBBX17* are gibberellin responsive (this work and Chu et al., 2016; Lira et al., 2020).

During tomato fruit development, GAs present a bimodal accumulation pattern (Gillaspy et al., 1993) that reflects their contribution in different phases of this process. A high GA level in flowers promotes fruit set and in young fruits induces growth through cell expansion, while GA level decreases during fruit ripening (Li et al., 2019) due to downregulation of GA-biosynthetic genes, such as *GA3ox1* and *GA20ox2*, and upregulation of GA deactivation genes, such as *GA2ox* (Zouine et al., 2017; Li et al., 2019). Consistently, exogenous application of GA_3_ to mature green fruit delayed fruit ripening (Li et al., 2019). In *Sl*BBX16OE and *Sl*BBX17OE fruits at the immature green stage, the modulation of genes regulating GA levels is not accompanied by a decline in bioactive GA content. However, in *Sl*BBX17OE fruits we observed an increased content of bioactive GAs, that can induce a homeostatic response in GA metabolism genes. The levels of bioactive GAs in plants are maintained via a complex regulation of GA metabolism (Hedden and Phillips, 2000; Olszewski et al., 2002). GA-deficient mutants show up-regulation of biosynthetic genes, GA20ox and GA3ox, and inhibition of the deactivation gene, GA2ox (Yamaguchi, 2008). On the other hand, GA treatment induces GA2ox and inhibits GA20ox and GA3ox (Yamaguchi, 2008). GA homeostasis is also controlled by GA signaling pathway, involving DELLA proteins, negative regulators degraded in the presence of GA. It has been demonstrated that DELLA regulates the expression of GA20ox2 by forming a complex with the transcription factor GAI-ASSOCIATED FACTOR 1 (GAF1) (Fukazawa et al., 2014; Livne et al., 2015). Our data suggests that phenotypical changes caused by *Sl*BBX16 and *Sl*BBX17 overexpression on fruit development are for both proteins related to modifications of GA homeostasis. What was observed by analyzing GA metabolism in a specific temporal window (young fruits of 1 cm in diameter) can be a snapshot of the homeostatic process in action (Figure 8).

**Figure 8.**
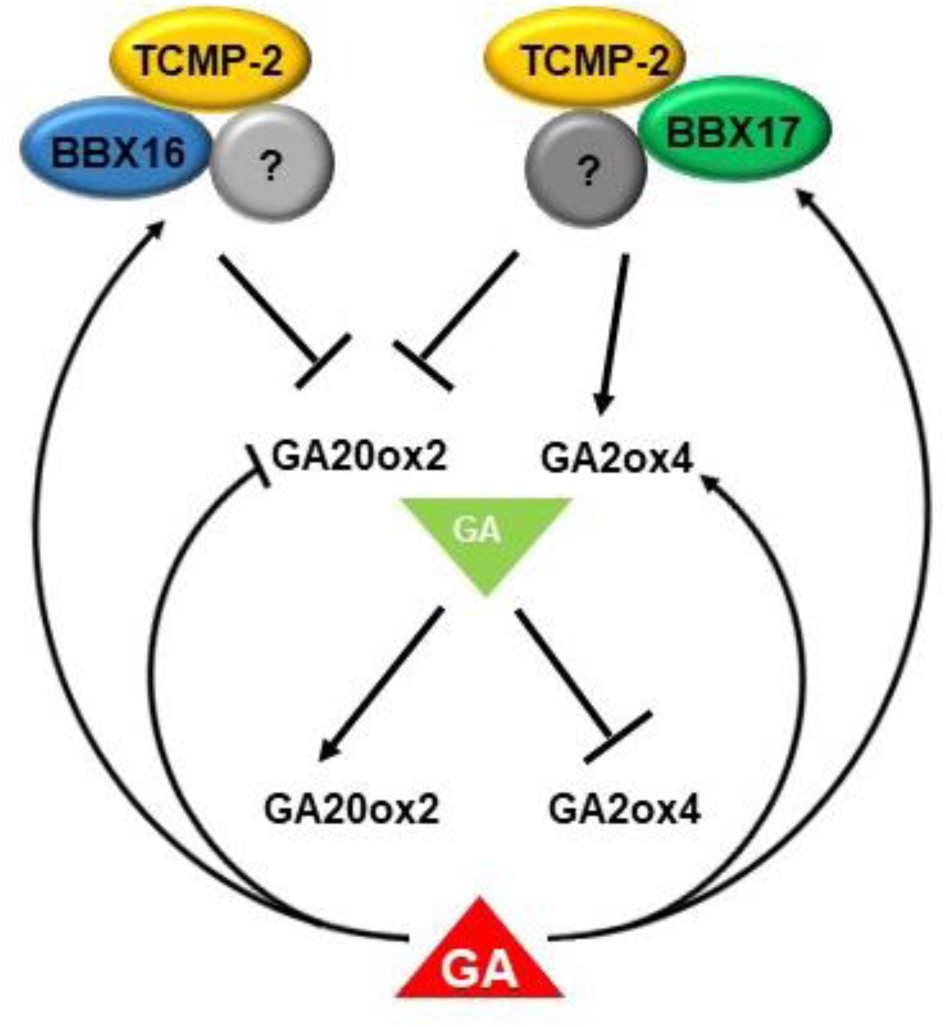
Interplay between *Sl*BBX16/17 and GA metabolism during fruit development. *Sl*BBX16 *and Sl*BBX17, which form regulatory complexes with TCMP-2 and presumably other unknown factors (?), when overexpressed cause inhibition of GA20ox2 and induction of GA2oX4, alterations ascribable to high GA level or signalling. These transcriptional changes would result in feedback regulation of GA metabolism and *SlBBX16/17* transcription. Green triangle= low bioactive GAs; red triangle= high bioactive GAs.

It will be critical for future research to decipher the regulatory functions of these tomato microProteins in the reproductive process, investigating whether they may be part of a multiprotein complex together with TCMP-2. Ultimately, a better understanding of the mechanism of action by which these microProteins alter GA homeostasis will provide opportunities for their potential application in tomato breeding to match the reproductive phases with changes in climate conditions.

## Materials and Methods

### Plant materials

*Solanum lycopersicum* MicroTom WT seeds (ID:TOMJPF00001) were obtained from the TOMATOMA mutant archive (Saito et al., 2011). *Arabidopsis thaliana* WT (ecotype Col-0) and *At*miP1a/b double KO mutant plants were employed for flowering time assessment.

### Plant genetic transformation

The tomato DNA sequences corresponding to the coding regions of *SlBBX16* (Solyc12g005750) and *SlBBX17* (Solyc07g052620) were amplified by PCR from cDNAs using the primers reported in Table S1. The DNA fragments were subcloned into the pGEM®-T Easy Vector (Promega) and checked by sequencing. The coding regions of *SlBBX16* and *SlBBX17* were then cloned into a derivative of the pBin19 vector, under the control of the *CaMV35S* promoter and the terminator sequence of the *Agrobacterium tumefaciens* nopaline synthase gene.

For the overexpression of *TCMP-2* (Solyc07g049140) in MicroTom, a sequence corresponding to the coding region was amplified using the Gateway System (Invitrogen) (Table S1). After subcloning in the pDONR221, the resulting pENTRY vector was checked by sequencing and used for recombination in the destination vector pK7WG2D.1 (Karimi et al., 2002). The recombinant vectors obtained were introduced into *Agrobacterium tumefaciens* cells (strain GV2260). The genetic transformation of MicroTom was obtained from cotyledon explants of 8 days-old seedlings. *A. tumefaciens* cells were grown at 28°C for 24 hours and used at OD_600_ of 0.1 for explant infection. After 48 hours of co-cultivation, the explants were transferred to culture medium containing NAA (0.01 mgL^−1^), zeatin riboside (2 mgL^−1^) and kanamycin (100 mgL^−1^). The regenerated shoots were transferred to rooting medium supplemented with kanamycin (75 mgL^−1^). After 3-4 weeks, the rooted plants were acclimatized in the greenhouse. Monitoring the ploidy level of putative transformants according to Atarés et al., 2011, we demonstrated that with this transformation method, the vast majority (about 90%) of MicroTom transformed lines retained the diploid state. *Arabidopsis* plants were transformed using the floral dip method (Zhang et al., 2006).

### Phenotypic analysis

Tomato plants were grown in the greenhouse during springtime. For phenotypic assessment, plants of the T1 generation were grown in pots and transgenic state was confirmed by spraying with kanamycin (400 mgL^−1^). Various flowering and fruiting parameters were recorded, and the fruit yield was evaluated at about 110 days after sowing. *Arabidopsis* plants were grown in a climatic chamber at a constant temperature of 25°C under LD conditions (16/8 hours light/dark cycle, photosynthetic photon fluence rate of 150 μmol m*^−2^* s^−1^). Homozygous plants of the T3 generation were used for flowering time analysis.

### Yeast Two-Hybrid analysis

To examine protein-protein interactions, the Matchmaker Gold Yeast Two-Hybrid System (Clontech) was used, following the manufacturer’s instructions with minor modifications. To test the interaction between TCMP-2 and *Sl*BBX17, the DNA sequence of the mature portion of the TCMP-2 protein (Solyc07g049140; from amino acid 53 to 96) was expressed as a fusion with the DNA-binding domain of GAL4 in the pGBKT7-BD vector, while the entire coding region of *SlBBX17* was cloned in frame into pGADT7-AD. For the interactions between *Sl*BBX16, *Sl*BBX17, *At*CO (*At*5g15840) and *Sl*COL1 (Solyc02g089540), the entire coding regions of the BBX genes were cloned in frame into the pGBKT7-BD vector and *At*CO and *Sl*COL1 were cloned in the pGADT7-AD vector. The interaction between miP1a (At3g21890) and *At*CO represents the positive control. For negative controls, pGBKT7 without insert (BD alone; Empty) and pGADT7 without insert (AD alone; Empty) were used. The primers employed for the genetic constructs preparation are reported in Table S1.

### RT-qPCR analysis

Total RNA extraction was performed from leaves, floral organs and fruits using the “NucleoSpin RNA Plant” kit (Macherey-Nagel). After DNase I treatment, first-strand cDNA was synthesized using the ImProm-II Reverse Transcriptase (Promega). Three cDNA samples derived from three independent RNA extractions were synthesized and amplified using Luna^®^Universal qPCR Master Mix (New England Biolabs, Ipswich, MA, USA) on a QuantStudio 3 Real-Time PCR System (Thermo Fisher Scientific). The data were normalized using *a*ctin (Solyc11g005330) or *SAND* (Solyc03g115810) as references for leaf and fruit tomato samples, respectively, and actin (At3g18780) for genes expressed in *Arabidopsis*. Data analysis was performed using the 2^−ΔΔCt^ method (Livak and Schmittgen, 2001).

### GA treatment and quantification

To test the responsiveness of *SlBBX16* and *SlBBX17* to the exogenous application of GA (GA_3_), tomato seedlings were grown *in vitro* for one month on half-strength MS agar medium (pH 5.9). Plants were transferred in a half-strength MS liquid solution (pH 5.9) and the next day, treated for 24 h with 5 µM GA_3_. The shoots were collected for expression analysis by RT-qPCR.

GAs extractions and quantifications from tomato immature green fruits were conducted at the Instituto de Biología Molecular y Celular de Plantas, Universidad Politécnica de Valencia using Q-Exactive mass spectrometer coupled to Ultra HighPerformance Liquid Chromatography. Each biological replicate was obtained by pooling 3-5 immature green fruits collected from at least four individual plants.

### Statistical analysis

Statistical analyses were performed using GraphPad Prism version 5.0 software (GraphPad Software, San Diego California USA). Data were compared using Student’s t-test.

## Acknowledgements

This research was supported by a MIUR grant (PRIN2017, number 20173LBZM2) given to T.P. University of Verona, Italy. The laboratory of SW receives funding through NovoCrops Centre (Novo Nordisk Foundation project number 2019OC53580), the Independent Research Fund Denmark (0136-00015B and 0135-00014B) and the Novo Nordisk Foundation (NNF18OC0034226 and NNF20OC0061440).

## Author contributions

A. M., and T.P. conceived and designed this work; V.D., F.P., and D.F carried out the molecular analyses and the genetic transformation of *Arabidopsis* and tomato plants; A.A contributed to the analysis of the transformed plants; S.W. provided *Arabidopsis* mutant; B.M., S.W., and T.P. discussed the results, B.M. and T.P. wrote the manuscript. All authors read and approved the final manuscript.

## SUPPORTING INFORMATION

**Figure S1.**
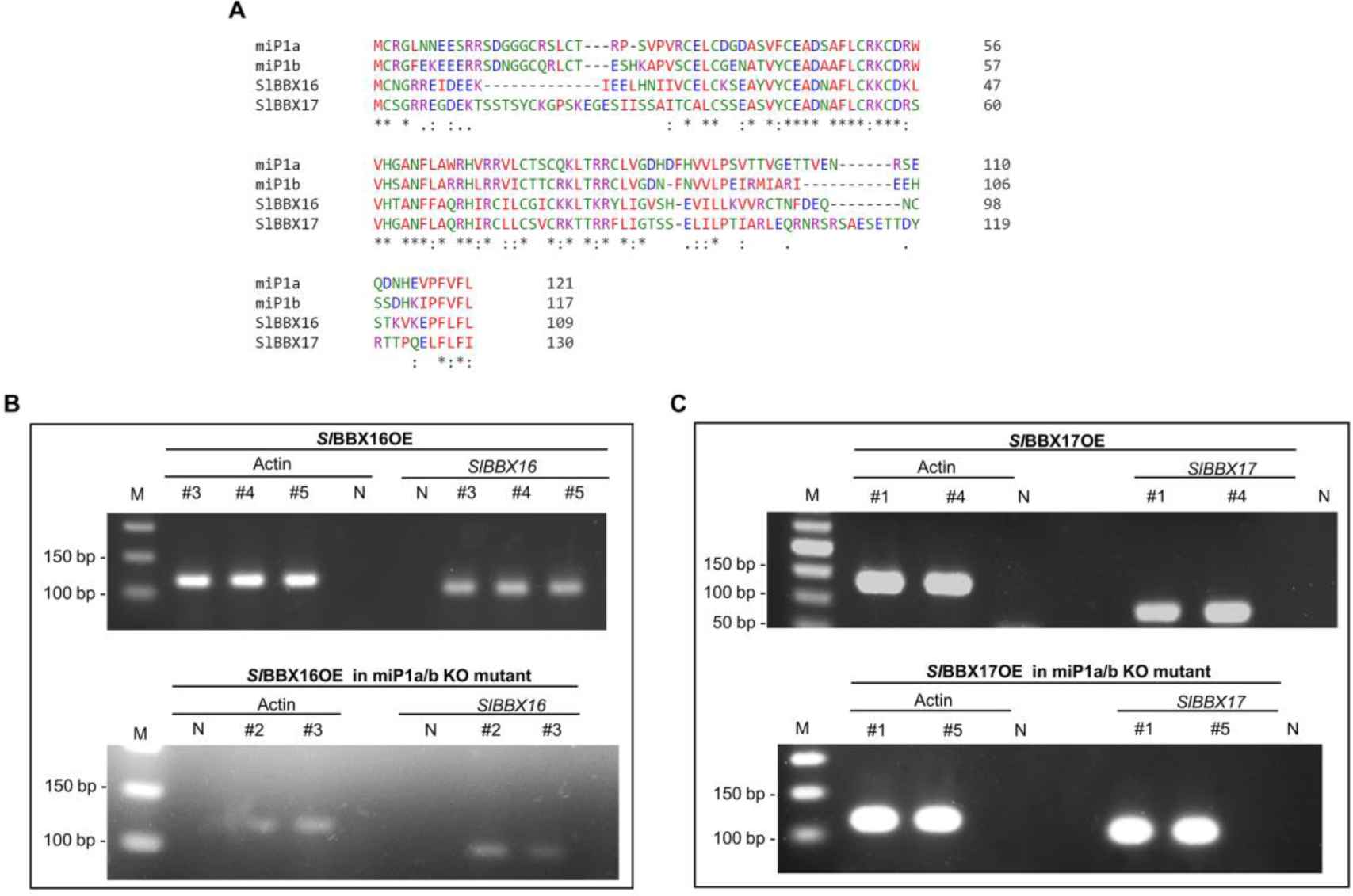
*SlBBX16* and *SlBBX17* overexpressing lines in *Arabidopsis*. **(A)** Sequence alignment of *Sl*BBX16 and *Sl*BBX17 with the closest homologs in *Arabidopsis*, miP1a and miP1b. The amino acid sequences were aligned using the ClustalW2 (http://www.ebi.ac.uk/Tools/msa/clustalw2) program, choosing default alignment parameters and selecting BLOSUM for the protein matrix. Consensus symbols: ‘*’ identical residues; ‘:’ residues with strongly similar properties; ‘.’ residues with weakly similar properties. **(B, C)** RT-PCR analysis carried out on *Sl*BBX16OE lines and *Sl*BBX17OE lines both in *Arabidopsis* WT and miP1a/b double KO mutant. N= no template control; M= DNA ladder.

**Figure S2.**
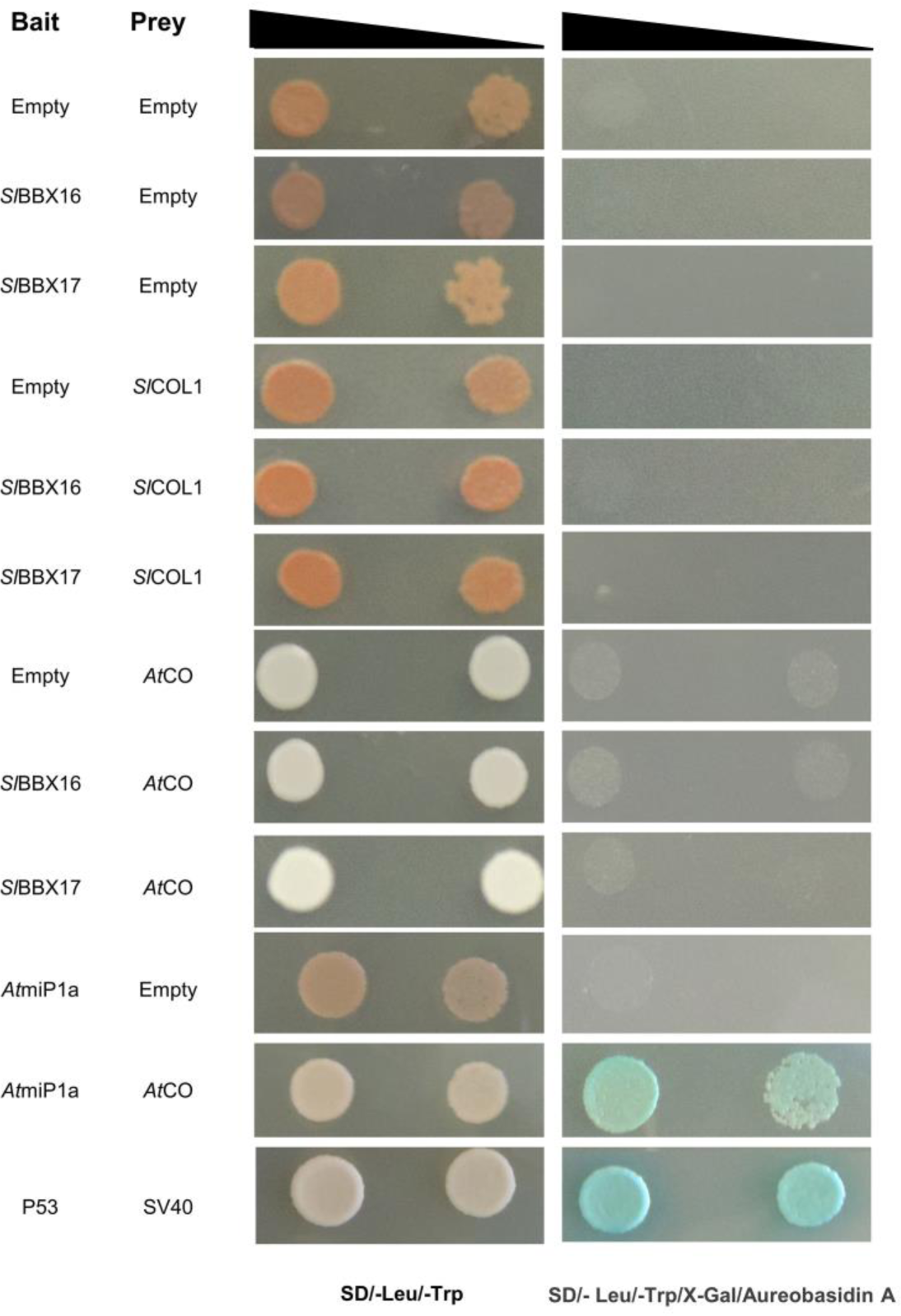
Y2H interaction assay of *Sl*BBX16 and *Sl*BBX17 with *At*CO and *Sl*COL1. The empty bait and prey vectors (BD and AD) were used to detect self-activation. For each protein-protein interaction, two increasing dilutions of the mated cultures (10^−2^ and 10^−3^) were spotted on control medium (SD/-Leu/-Trp) and selection medium plates (SD/-Leu/-Trp/X-Gal/Aureobasidin A). The interaction of *At*miP1a with *At*CO (Graeff et al., 2016) and p53 with SV40 were used as positive mating controls.

**Figure S3.**
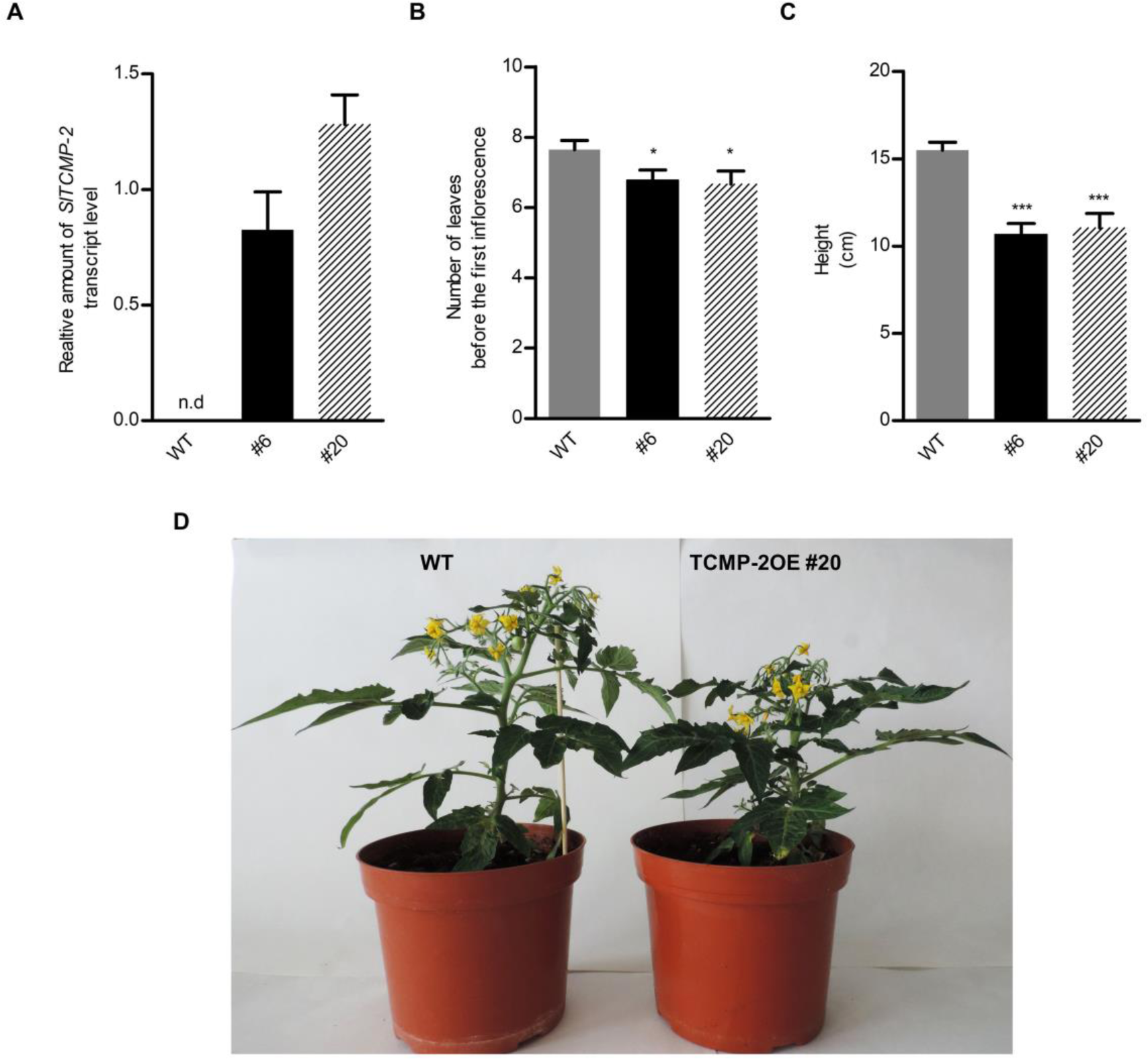
Overexpression of *TCMP-2* in the cultivar MicroTom. **(A)** Expression level of *TCMP-2* evaluated by RT-qPCR in tomato leaves of WT and two overexpressing lines (#6 and #20). Values are mean ± SE of three biological replicates; n.d. means no detectable. Line #20 was used as calibrator. **(B)** Number of leaves before the first inflorescence, **(C)** total plant height. Values are mean ± SE (n=9-10 for panels B and C). **(D)** Phenotypical aspect of WT and TCMP-2OE #20 plants at the same developmental stage.

**Figure S4.**
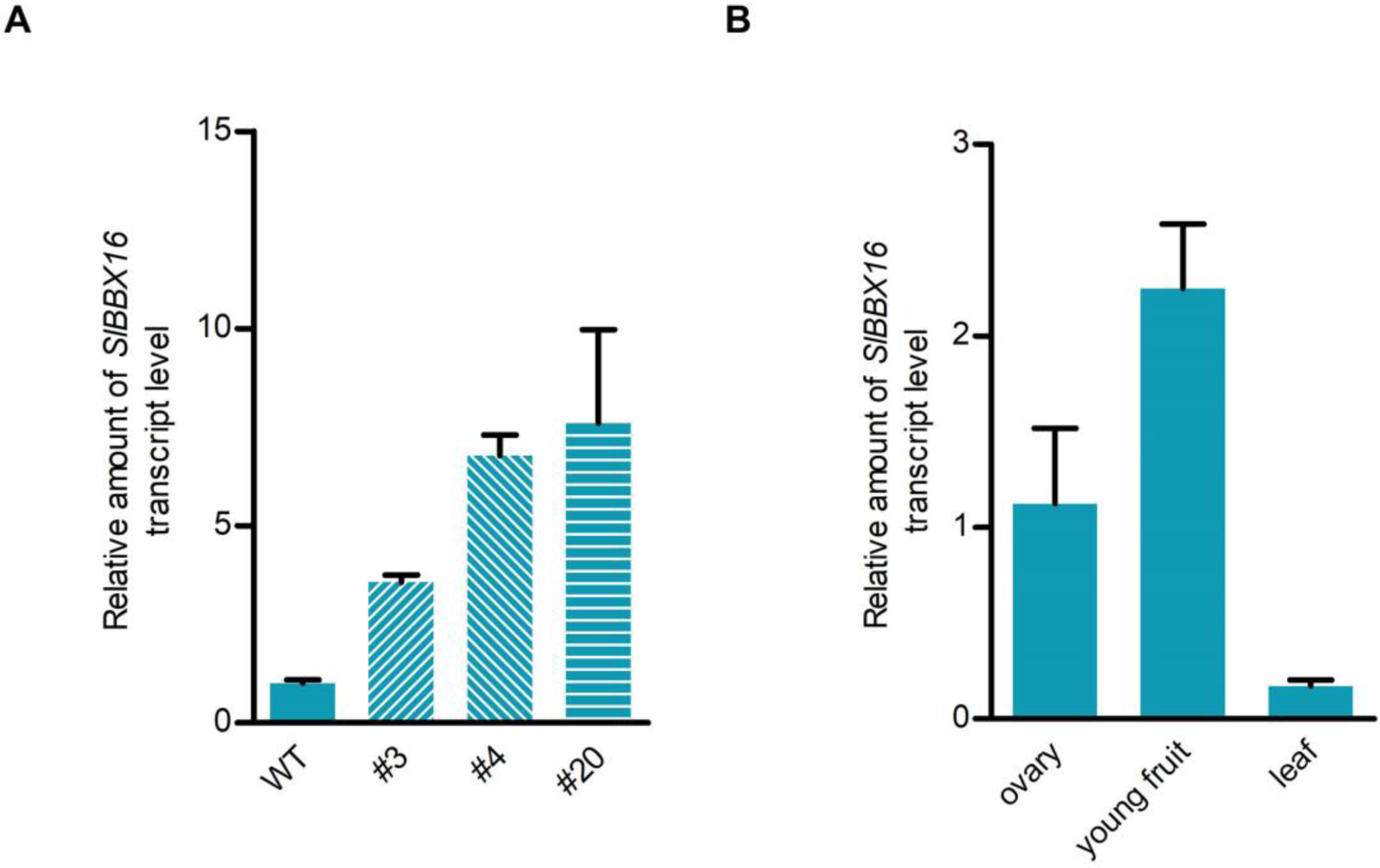
Tomato *SlBBX16* overexpressing lines and expression of *SlBBX16* in different organs. **(A)** *SlBBX16* expression in three transgenic lines (#3, #4, and #20). WT was used as calibrator. **(B)** Expression level of *SlBBX16* in ovary detached from flowers at anthesis, young fruits (0.5-1.0 cm diameter) and leaves. Ovary was used as calibrator. Values are mean ± SE of three biological replicates.

**Figure S5.**
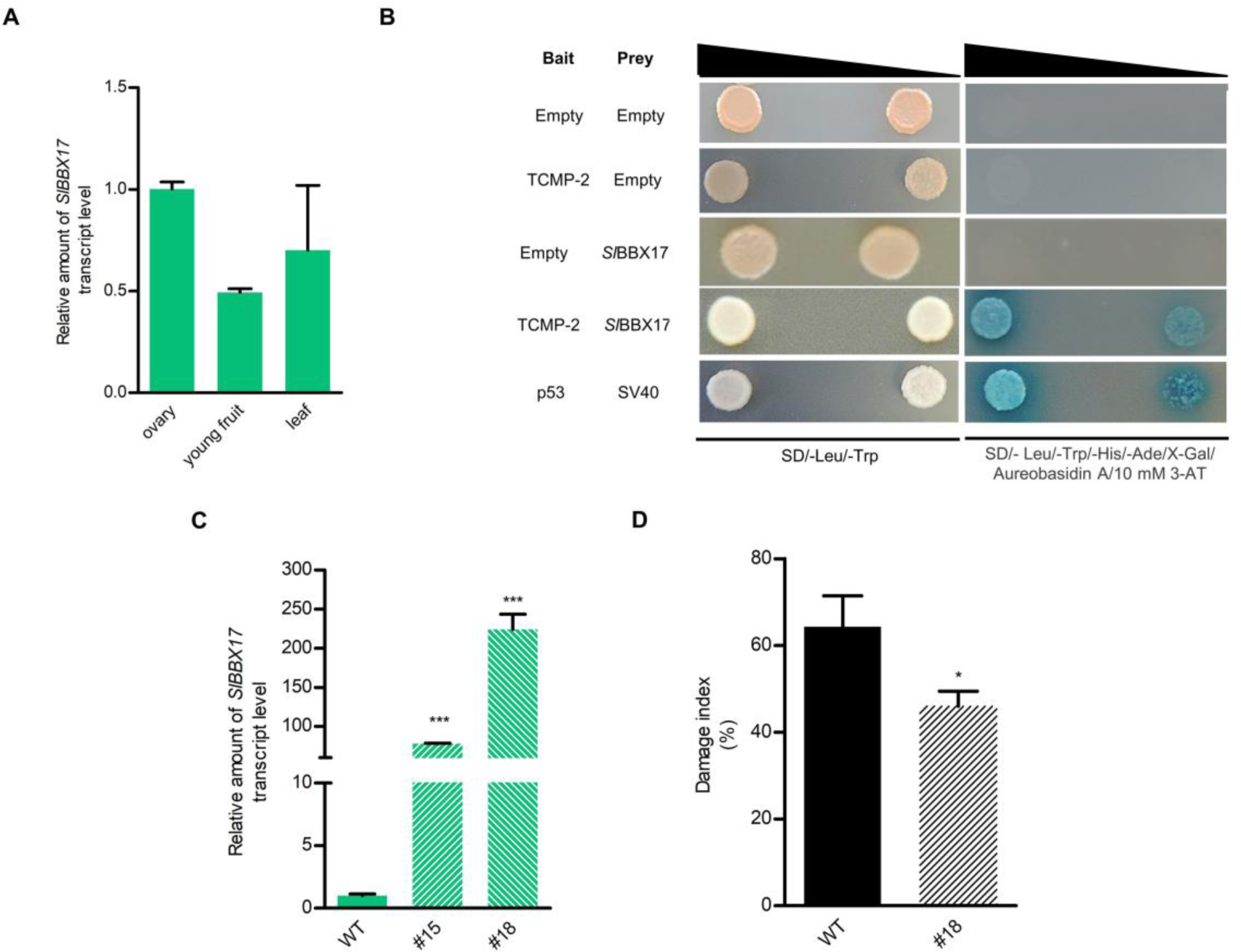
**(A)** Expression level of *SlBBX17* in ovary detached from flowers at anthesis, young fruits (0.5-1.0 cm diameter) and leaves. Ovary was used as calibrator. Values reported are means ± SE of three biological replicates. **(B)** Y2H interaction assay between TCMP-2 and *Sl*BBX17. For negative controls, pGBKT7 without insert (BD alone; Empty), pGADT7 without insert (AD alone; Empty) were used. For each protein-protein interaction, two increasing dilutions of the mated cultures (10^−2^ and 10^−3^) were spotted on control medium (SD/-Leu/-Trp) and selection medium plates (SD/-Leu/-Trp/-His/-Ade/X-Gal/Aureobasidin A/10 mM 3-AT). The interaction of p53 with SV40 was used as a positive mating control. **(C)** Expression of *SlBBX17* in tomato transgenic lines (#15 and #18). WT was used as calibrator. Values reported are means ± SE of three biological replicates. **(D)** Heat stress response of one-month-old WT and line #18 plants. Damage index was calculated as the percentage of dead or severely damaged plants after imposing heat stress at 42°C for 24 h. Values are mean ± SE of three independent experiments. *P < 0.05, ***P < 0.001 versus the respective control (Student’s t-test).

**Figure S6.**
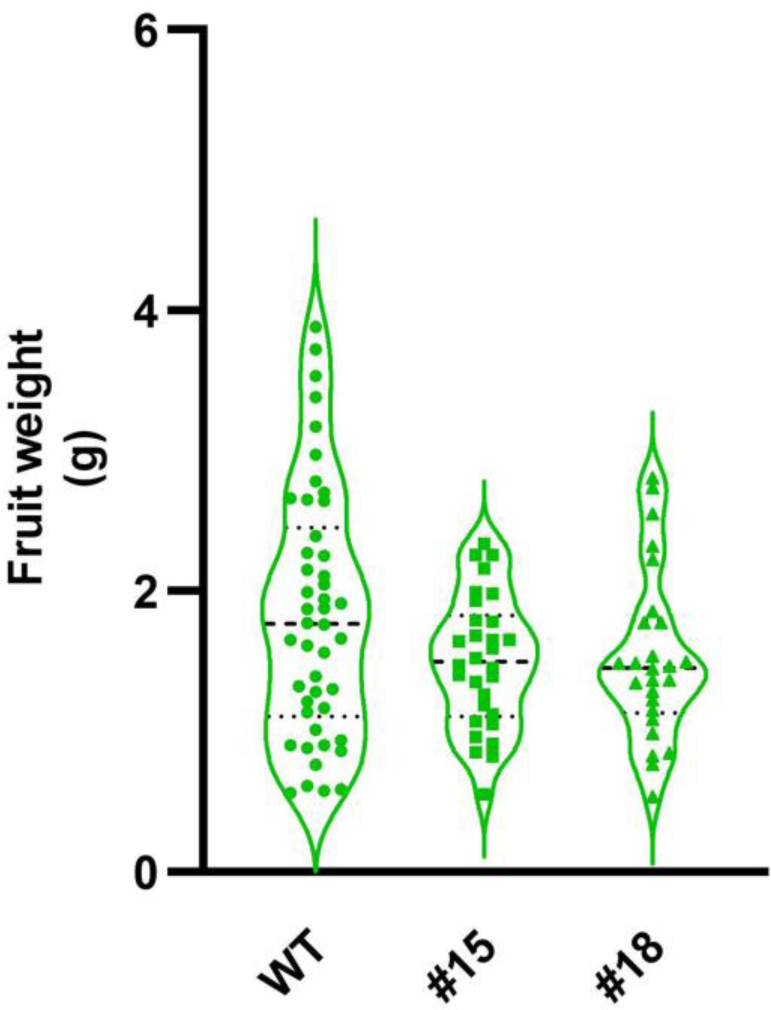
Green fruit weight of *Sl*BBX17OE. Comparison between distribution of green fruits’ weights in WT and *Sl*BBX17OE lines. Dashed lines represent median and dotted lines quartiles.

**Table S1.**
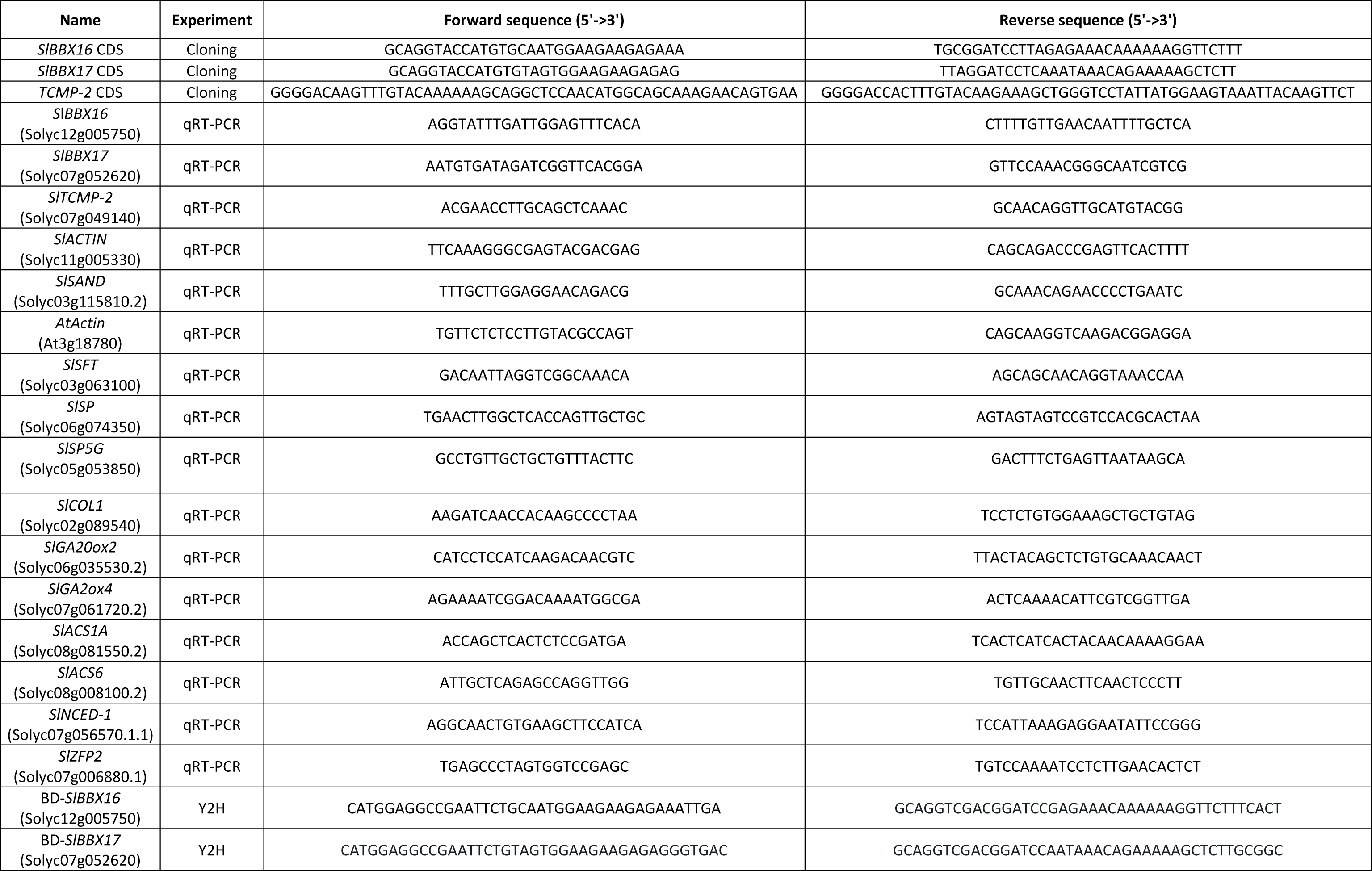
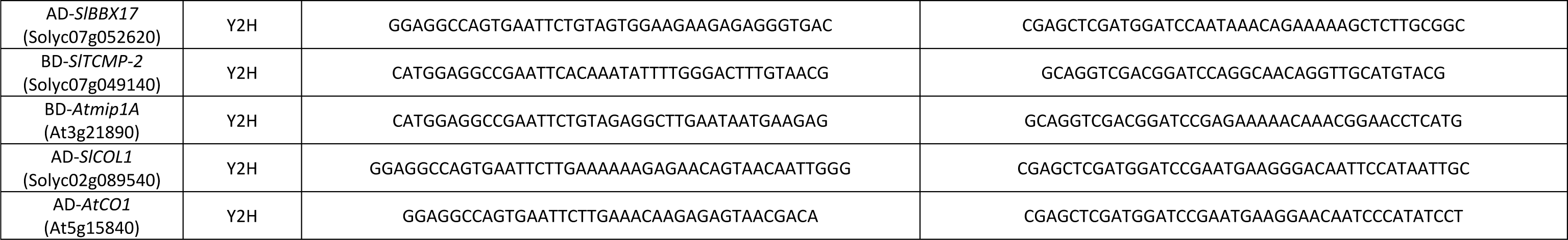
List of primers.

